# Reconstitution of an N-AChR from *Brugia malayi*

**DOI:** 10.1101/2022.06.01.493574

**Authors:** Jennifer D. Noonan, Robin N. Beech

## Abstract

Neurotransmission is an important target for anthelmintic drugs, where receptor characteristics and response can be examined through reconstitution *ex vivo* in *Xenopus laevis* oocytes. The homomeric ACR-16 nicotine sensitive acetylcholine receptors (N-AChRs) of several helminth species have been characterized in this way. Our efforts to reconstitute the N-AChR from the clade III filarial parasite, *Brugia malayi* using similar conditions, initially produced no detectable response. A robust response to acetylcholine is obtained from the closely related clade III parasite *Ascaris suum*, suggesting that specific changes have occurred between *Ascaris* and *Brugia*. N-AChRs from three species intermediate between *A. suum* and *B. malayi* were characterized to provide information on the cause. Maximal current to acetylcholine did not change abruptly, consistent with a discrete event, but rather decreased progressively from *A. suum* through *Dracunculus medinensis*, *Gonglylonema pulchrum* and *Thelazia callipaeda.* Receptor responses to the characteristic nicotine, and other agonists were generally similar. The decrease in maximal current did correlate with a delayed time to maximal response. Together, this suggested that the failure to reconstitute the *B. malayi* N-AChR was one extreme of a progressive decrease and that synthesis of the receptor in oocytes was responsible. Addition of accessory proteins EMC-6, NRA-2 and NRA-4, in addition to RIC-3, produced a small, but measurable *B. malayi* N-AChR response. Pharmacological properties of a chimeric *B. malayi* N-AChR were equivalent to the other species, confirming the receptor response remains unchanged while its production is increasingly dependent on accessory proteins. One possibility is that loss of many subunits for acetylcholine receptors from the filarial nematode genome is linked to such a dependence. This novel phylogenetic approach allowed the first characterization of a *B. malayi* AChR *ex vivo* and in doing so, provides a framework for the successful characterization of other receptors that have yet to be reconstituted.

## Introduction

Random drugs screens for antiparasitic compounds have produced several that target neurotransmission and suggest post-synaptic, neurotransmitter receptors are useful targets for anthelmintic drugs [1–4]. These pentameric ligand-gated ion channels (pLGICs) are composed of five subunits that form a radially symmetrical complex embedded in the post-synaptic membrane [5]. Subunits of these channels are encoded by a gene family, derived from an ancestral gene shared with the prokaryotes [6]. This family has expanded in the nematodes and is characterized by many gene duplication and loss events [7,8]. Their 3D structure is highly conserved where each subunit contains a large extracellular domain (ECD) [9], four transmembrane domains (TM 1-4) [10] and a large, variable intracellular loop (ICL) [11]. Activating ligands bind at the ECD interface between adjacent subunits. Anthelmintics that activate these channels lead to paralysis, death and/or expulsion of the worm [12–14] and so represent a significant biological target for the control of infection [15].

Identification of different target receptors has typically been achieved using genetic techniques in the model nematode *Caenorhabditis elegans* [3,14,16–19]. Confirmation of the composition and characteristics of specific receptors has then been facilitated by expression of subunits, *ex vivo*, in the oocytes of *Xenopus laevis* [1,2,20–22]. More recently, identification of predicted subunit genes in genome data, followed by direct expression in oocytes has proved an effective tool to identify previously unknown receptor classes directly [2]. There are limitations to this technique since expression of receptors *ex vivo* occurs in the absence of the many accessory proteins they normally interact with during assembly and stabilization of precursors to the mature receptor complex *in vivo* [23]. More than a dozen accessory proteins have been characterized in *C. elegans* that interact with the acetylcholine receptors (AChRs) although only a few are required *ex vivo*. These include RIC-3, that is required for the nicotine sensitive acetylcholine receptor (N-AChR) encoded by *acr-16* [1,24], and UNC-50 and UNC-74, together with RIC-3 that are required for the levamisole sensitive receptor (L-AChR) [1]. These proteins have homologs in vertebrates but their expression in the oocyte appears insufficient for reconstitution of the nematode receptors since over-expression of the *Xenopus* RIC-3 can successfully reconstitute the *C. elegans* N-AChR [25,26].

The filarial nematodes, such as *Brugia malayi*, are an important group of parasites where the range of antiparasitic drugs is limited and where the mechanism of action for drugs that are available is much less clear than for species more closely related to *C. elegans* [27]. Characterization of neurotransmitter receptors in filarial worms *ex vivo* will therefore be an important resource. Two acetylcholine-sensitive neurotransmitter receptors in the muscle regulate body movement in *C. elegans*, the N-AChR and L-AChR [28]. In parasitic nematodes, a third is present, encoded by *acr-26* and *acr-27*, that is sensitive to morantel (M-AChR) [2]. While four different acetylcholine receptors have been demonstrated in living *B. malayi,* representing the L-AChR, N-AChR, M-AChR and one sensitive to pyrantel (P-AChR) [29–31], none have been characterized in recombinant systems.

The N-AChR, a homomeric receptor composed of ACR-16, has been characterized in many different nematode species including *Ancylostoma caninum* [32]*, A. ceylanicum* [33]*, Necator americanus* [33]*, Trichuris suis* [34]*, T. muris* [35] and two species basal to the filarial nematode clade: *Ascaris suum* (Asu-ACR-16) [36] and *Parascaris equoram* (16). Unfortunately, initial attempts in our hands to reconstitute the *B. malayi* N-AChR (Bma-ACR-16) in the presence of RIC-3 were unsuccessful. To understand the reason for this, we characterized the N-AChR from three clade III nematode species intermediate between *A. suum* and *B. malayi*, namely *Dracunculus medinensis* (Dme-ACR-16), *Gonglyonema pulchrum* (Gpu-ACR-16) and *Thelazia callipaeda* (Tzc-ACR-16) [38]. Each one of these intermediates are independent representatives of N-AChR change along the clade III phylogeny. Using this approach, we were able to show that the response of the N-AChR from these species, to a variety of agonists was not substantially different from each other. The maximal response however, decreased and took longer to achieve progressively from *A. suum* to *T. callipaeda*, with no response from *B. malayi*. This suggested a progressive change along the phylogeny in receptor assembly and/or surface transport was responsible. Addition of accessory proteins EMC-6 [39], NRA-2 and NRA-4 [23,40] allowed reconstitution and characterization of the first *B. malayi* AChR *ex vivo*. The use of an evolutionary approach to follow the change in N-AChR function was a novel and effective way to investigate the mechanism responsible for the failure to reconstitute an N-AChR from *B. malayi*. This method may prove useful for the study of other helminth AChRs as drug targets.

## Results

### ACR-16 is conserved across nematodes

An N-AChR has been reconstituted and characterized as a homopentamer of ACR-16 from several different nematode species [24,32–34,36,37]. In each case, co-expression of the RIC-3 accessory protein is required for a robust response. These include the clade IIIb nematode, *A. suum* that encodes an N-AChR substantially similar to that found in the clade V nematode *C. elegans* [1,24,36]. The genome of the clade IIIc filarial parasite, *B. malayi* [41], also contains an *acr-16* that is expected to encode a functional N-AChR since the worm responds to nicotine (NIC) *in vivo* with an influx of cations as expected from functional AChRs [29,42]. Perhaps surprisingly, the *B. malayi* N-AChR failed to reconstitute a functional homomeric receptor *ex vivo* in the presence of RIC-3. Two likely explanations for this are either reconstitution of a receptor whose functional characteristics have changed sufficiently for the receptor to no longer respond to a panel of conventional ligands or that no functional receptor is produced in the oocyte expression system. The fact that the *A. suum* N-AChR can be reconstituted [36], but not *B. malayi* indicates a specific change has occurred in more derived clade III ACR-16. To identify this change and when it occurred along filarial phylogeny, we characterized the N-AChR from three intermediate species (Fig 1).

**Fig 1.**
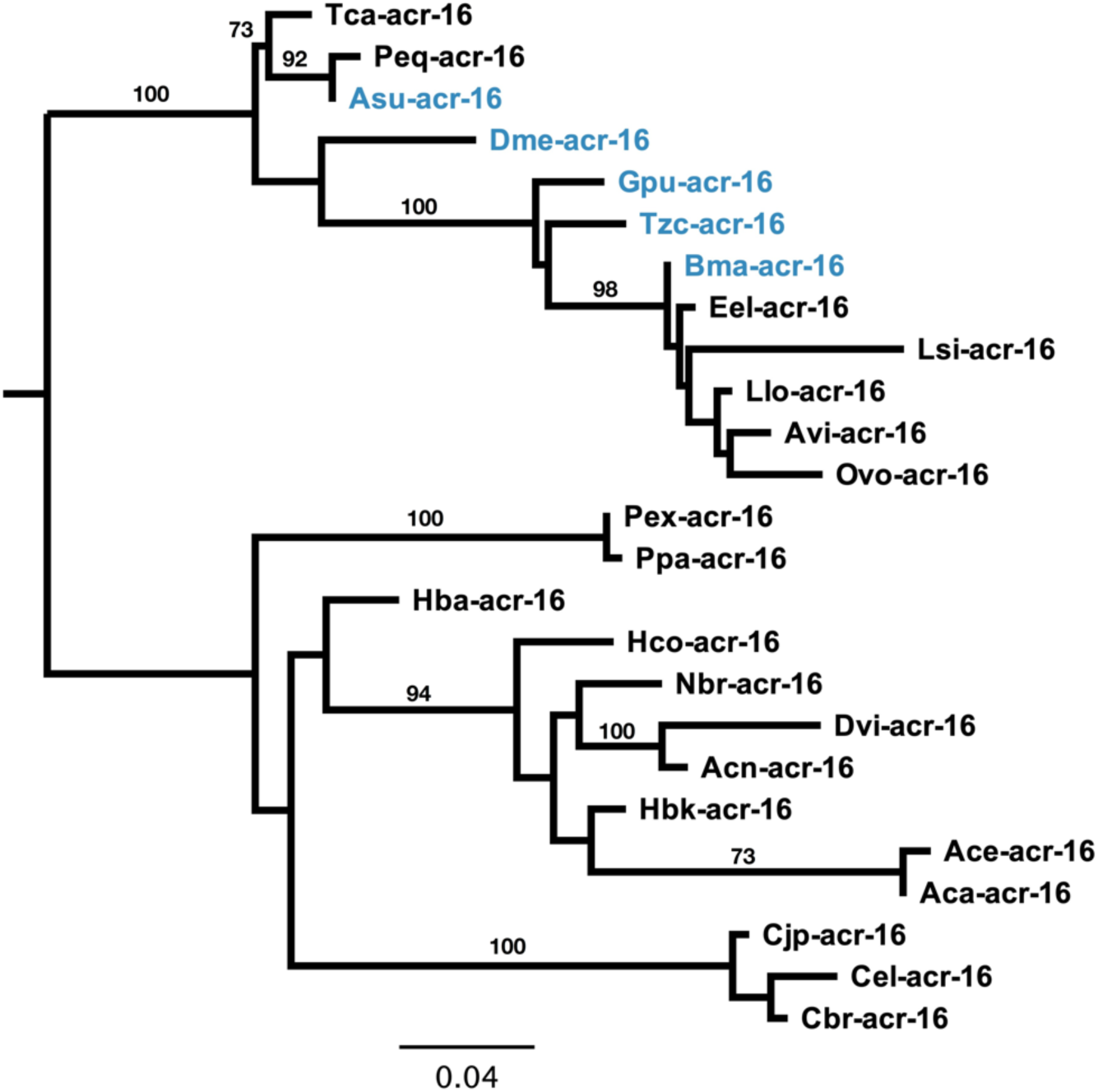
ACR-16 phylogeny in the nematodes. ACR-16 is conserved across worms with short branches observed within the clade III worms. Clade III and clade V ACR-16 group into their respective clusters with no unexpected branching topology of *B. malayi* ACR-16 would explain the inability to measure a *B. malayi* N-AChR receptor *ex vivo*. ACR-16 sequences used in this study indicated in blue. Bootstrap resampling branch support values are shown.

The phylogeny and aligned primary sequence of ACR-16 for *C. elegans*, *A. suum*, *D. medinensis, A. G. pulchrum, T. callipaeda* and *B. malayi* in order of divergence from *A. suum* are shown in Fig 1 and Fig 2. A majority of changes occurred within the intracellular loop, as expected since this region is commonly found to be the most variable region within pLGIC receptors [11]. The transmembrane domains had few changes, confined to TM2 and TM4. The ion charge filter motif, located at the start of TM2, (GEK) was conserved throughout, as such receptors were expected to be cation permeable [10]. Substitution within TM2 is similar in side chain chemistry (I287V) and is therefore not expected to alter ion-selectivity. Substitutions within the six agonist binding loops were not residues that interact with cholinergic ligands, so it seems unlikely that a significant change in activating ligand has occurred [43,44]. Overall, no substitutions within functionally important regions were found within the *B. malayi* ACR-16 sequence that could explain the inability to reconstitute a detectable receptor in oocytes. We next wanted to determine the function of each N-AChR in oocytes.

**Fig 2.**
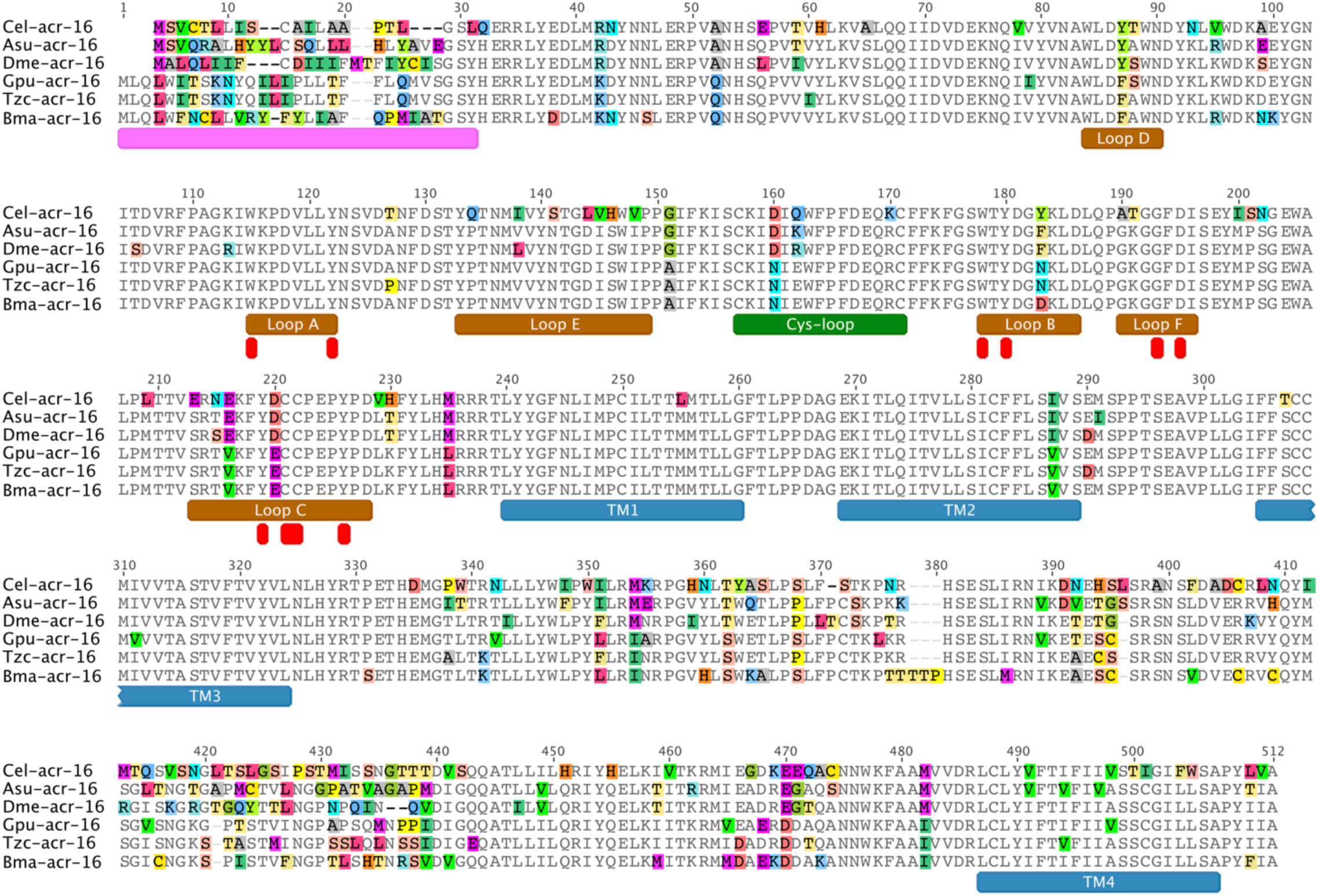
ACR-16 is conserved in clade III worms. Five clade III ACR-16 were used to identify why no *B. malayi* N-AChR can be measured in oocytes. Predicted coding sequences were synthesized from their respective genomes obtained from WormBase ParaSite WBPS13 [81,87] with primary protein sequence alignment shown. All functionally important regions and motifs are conserved within the *B. malayi* sequence and implies that no specific substitutions explain the inability of measuring an N-AChR. Transmembrane domains are shown in blue, signal peptide in pink, acetylcholine binding loops in brown, cys-loop in green and residues that interact with agonist in red [43,44]

### Clade III N-AChRs show progressive declining maximal response

Each ACR-16 subunit was co-injected with *H. contortus* RIC-3 and, except for *B. malayi*, produced measurable agonist-induced currents. A progressive decline in maximal response to both acetylcholine (ACh) and NIC was observed from *A. suum* to *B. malayi* (Fig 3A-B). Receptor currents were compared to Asu-ACR-16 as it lies at the base of the clade and has been previously characterized [36,38]. For acetylcholine-induced currents, Asu-ACR-16 (2052 ± 66 nA) and Dme-ACR-16 (2150 ± 59 nA, p=0.2787) produced the largest and did not differ statistically (Fig 2A). In phylogenetic order, Gpu-ACR-16 produced smaller currents (1401 ± 91 nA, p<0.0001) and Tzc-ACR-16 produced even smaller measurable currents (563 ± 39 nA, p<0.0001). Bma-ACR-16 produced no detectable response (0 ± 0 nA, p< 0.0001). For nicotine-induced currents, Asu-ACR-16 (2067 ± 74 nA) and Dme-ACR-16 (2025 ± 81 nA, p=0.7134) again produced the highest currents that were not statistically different. Gpu-ACR-16 produced significantly smaller currents (1231 ± 98 nA, p<0.001), and Tzc-ACR-16 produced the smallest measurable currents (481 ± 45 nA, p<0.0001). Bma-ACR-16 again produced no measurable current (0 ± 0 nA, p< 0.0001). The same decline in response was observed when *B. malayi* RIC-3 replace *H. contortus* RIC-3 and was therefore not caused by incompatibility with the clade V accessory protein (Fig 3C). The consistent gradual decline in ACR-16 current along its phylogeny ruled out a discrete evolutionary event as responsible and instead suggested a progressive change in the ability to reconstitute a receptor *ex vivo*. We next characterized each N-AChR to determine if ligand-response characteristics contribute to the observed declining currents.

**Fig 3.**
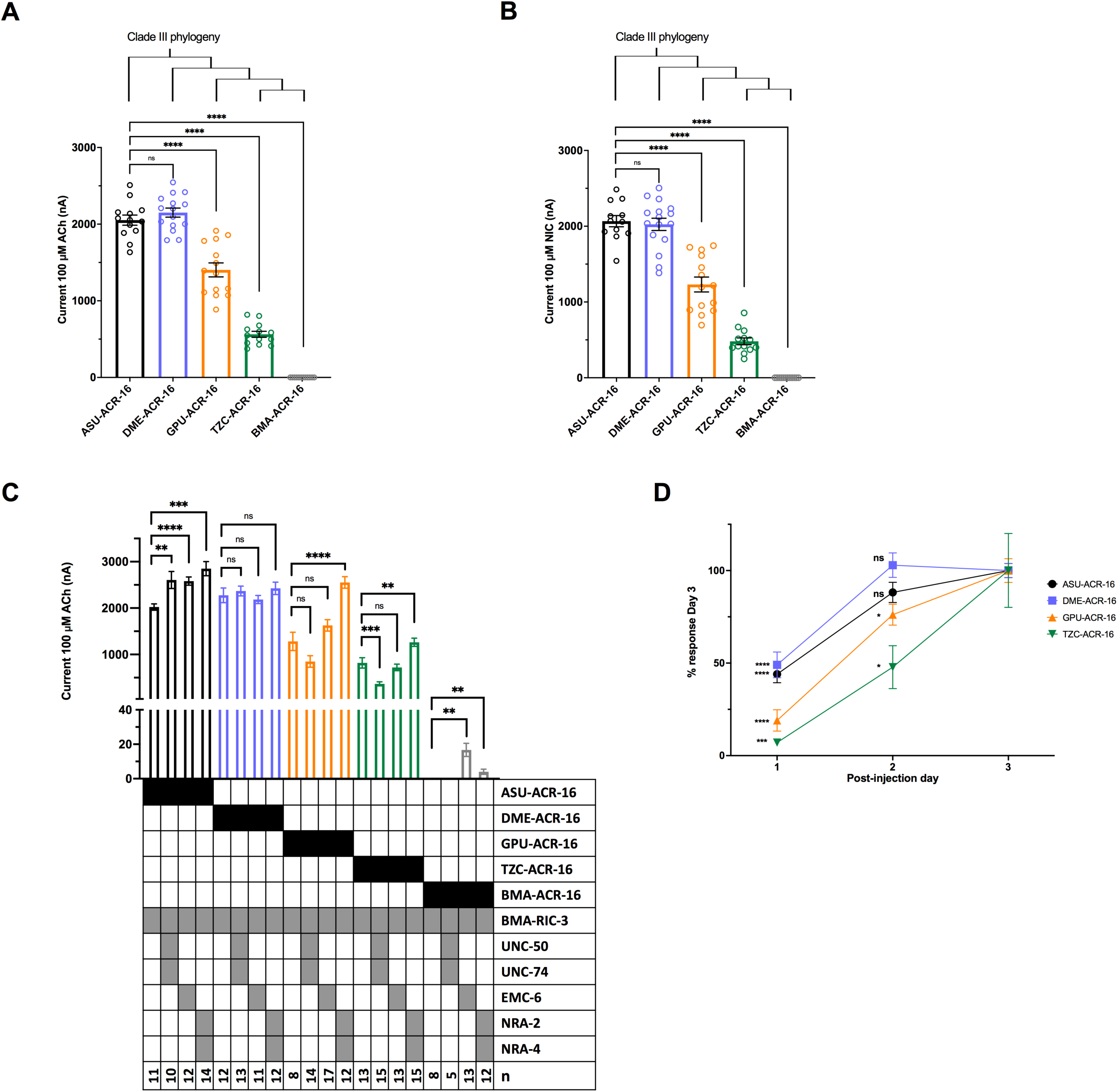
N-AChR has phylogenetic declining response and delayed expression in oocytes. To identify why no *B. malayi* N-AChR was measured in a heterologous expression system, N-AChRs from related species, along the clade III phylogeny, were reconstituted in oocytes. (A) A declining response to 100 µM acetylcholine was measured when co-injected with *H. contortus* RIC-3*. A. suum* served as the positive reference receptor. In order of phylogeny, responses to acetylcholine were 2052 ± 66 nA (n=13, black) for *A. suum;* 2150 ± 59 nA (n=15, p<0.0001, blue) for *D. medinensis;* 1403 ± 91 nA (n=14, p<0.0001, orange) for *G. pulchrum*; 563 ± 39 nA (n=13, p<0.0001, green) for *T. callipaeda*, 0 ± 0 nA (n=12, p<0.0001, grey) for *B. malayi*. ****p<0.0001. (B) Responses to 100 µM nicotine were 2067 ± 74 nA (n=12, black) for *A. suum;* 2025 ± 81 nA (n=15, p<0.0001, blue) for *D. medinensis;* 1231 ± 98 nA (n=14, p<0.0001, orange) for *G. pulchrum*; 481 ± 45 nA (n=13, p<0.0001, green) for *T. callipaeda*, 0 ± 0 nA (n=12, p<0.0001, grey) for *B. malayi*. The declining response indicates a progressive adaptation has occurred within the clade that decreases response to ligand. Error bars represent standard error, circles represent recordings from individual oocytes. ****p<0.0001. (C) A declining response to 100 µM acetylcholine was obtained with the addition of other accessory protein conditions: *B. malayi* RIC-3; *B. malayi* RIC-3 + UNC-50 + UNC-74; *B. malayi* RIC-3 + *C. elegans* EMC-6; and *B. malayi* RIC-3 + *C. elegans* NRA-2 + NRA-4. Small *B. malayi* N-AChR responses were measured when including EMC-6 (17 ± 4 nA, n=13, p<0.01) or NRA-2 + NRA-4 (4 ± 2 nA, n=12, p<0.01). Error bars represent standard error. See S1 Fig for additional accessory protein conditions and S1 Table for values. **p<0.01; ***p<0.0005; ****p<0.0001. (C) An expression analysis was carried out to measure the time taken to reach maximal response for each N-AChR. Responses to 100 µM acetylcholine was measured for three consecutive days post-injection and normalized to responses obtained on Day 3. In order of phylogeny, 44 ± 5 % (n=12, p<0.0001) and 88 ± 5% (n=10, p=0.0841 ) for *A. suum* in black, 49 ± 7 % (n=10, p<0.0001) and 102 ± 7 % (n=10, p=0.7011) for *D. medinensis* in blue, 19 ± 6 % (n=17, p<0.0001) and 76 ± 6 % (n=14, p<0.05) for *G. pulchrum* in orange, 7 ± 1 % (n=11, p<0.0005) and 47 ± 12 % (n=10, p<0.05) for *T. callipaeda* in green, for Days 1 and 2, respectively. *A. suum* and *D. medinensis* reached full response by Day 2 that is not significantly different from Day 3, in contrast to *G. pulchrum* and *T. callipaeda* responses on Day 2 that were significantly smaller. The delayed time to reach maximal responses confirms changes to receptor functional synthesis contribute to the declining response observed. Error bars represent standard error. *p<0.05; ***p<0.0005; ****p<0.0001.

### Clade III N-AChRs have similar pharmacological response in oocytes

Agonist affinity was measured with half maximal effective concentration (EC_50_) for both ACh (Fig 4A) and NIC (Fig 4B). Dose-response curves were comparable between species. Consistent with previous characterization of Asu-ACR-16 [36], EC_50_ for Asu-ACR-16 was 8.9 ± 0.3 μM for ACh and 5.1 ± 0.2 μM for NIC. EC_50_ for Dme-ACR-16 was 12.8 ± 0.4 μM and 3.8 ± 0.2 μM, for Gpu-ACR-16 was 10.5 ± 0.5 μM and 2.1 ± 0.1 μM, and for Tzc-ACR-16 was 17.4 ± 0.8 μM and 4.9 ± 0.4 μM, for ACh and NIC respectively. A higher affinity was observed for NIC compared to ACh for all N-AChRs, agreeing with the published characterization of Asu-ACR-16 [36] and all receptors reached saturation (example recordings are shown in Fig 4C-F).

**Fig 4.**
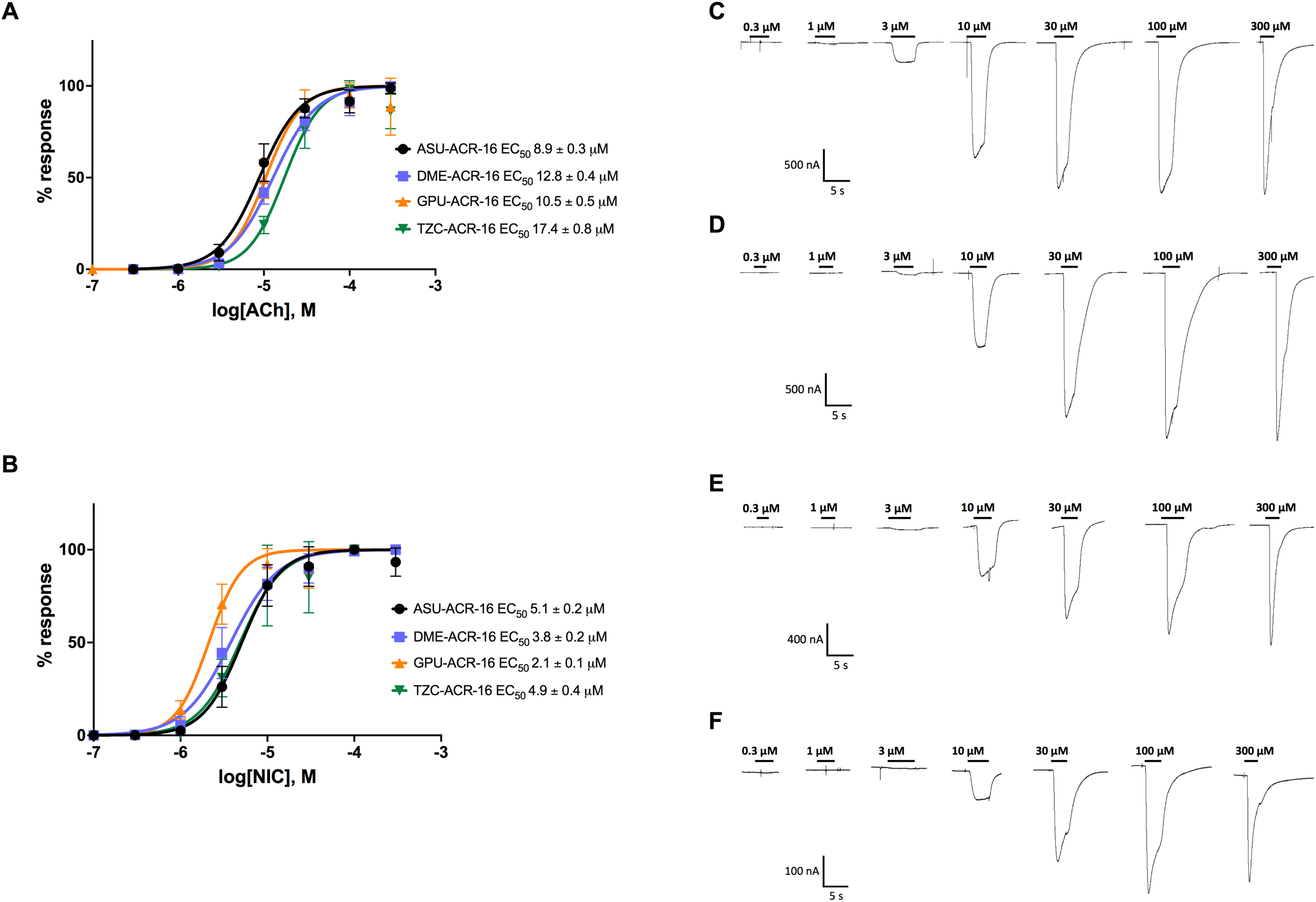
Clade III N-AChR show similar agonist affinities. Dose-response curves were measured to determine if declining maximal response is due to decreasing ligand affinity. Dose-response curves for clade III N-AChRs for (A) acetylcholine and (B) nicotine. In order of phylogeny; *A. suum* in black, *D. medinensis* in blue, *G. pulchrum* in orange, *T. callipaeda* in green. EC_50_ were *A. suum* 8.9 ± 0.3 µM (hill coefficient = 1.8 ± 0.1; n = 11) and 5.1 ± 0.2 µM (hill coefficient = 2.0 ± 0.1; n=13 ), *D. medinensis* 12.8 ± 0.4 µM (hill coefficient = 1.7 ± 0.1; n=11) and 3.8 ± 0.2 µM (hill coefficient = 1.6 ± 0.1; n=14), *G. pulchrum* 10.5 ± 0.5 µM (hill coefficient = 2.1 ± 0.2; n=15) and 2.1 ± 0.1 µM (hill coefficient = 2.3 ± 0.2; n=13), *T. callipaeda* 17.4 ± 0.8 µM (hill coefficient = 2.1 ± 0.2; n=10) and 4.9 ± 0.4 µM (hill coefficient = 1.9 ± 0.2; n=14), for acetylcholine and nicotine, respectively. Representative recordings in response to acetylcholine are shown for (C) *A. suum*. (D) *D. medinensis,* (E) *G. pulchrum* and (F) *T. callipaeda*. Similar dose-response curves confirm the declining response is not due to changes in agonist affinity. Error bars represent standard deviation.

Responses to each of the characteristic ligands for the four classes of AChRs identified in *B. malayi* were determined, namely levamisole (LEV), NIC, morantel (MOR), and pyrantel (PYR), [2,29,30,45,46]. Responses relative to ACh showed similar profiles across the species (Fig 5). Each receptor had a large response to NIC, smaller response to dimethylphenylpiperazinium (DMPP) and oxantel (OXA), and minimal response to MOR and pyrantel (PYR). No response was measured for LEV, epinephrine (EPI), norepinephrine (NOR), dopamine (DA), tyramine (TYR), betaine (BET) or bephenium (BEPH) for any receptor. Ligand-response profiles coincided with previous clade III N-AChR characterization [36,37] and confirm no meaningful change in ligand interaction has occurred. Taken together, the agonist-response panel and dose-response curves ruled out a pharmacological explanation for the trend observed in Fig 2A.

**Fig 5.**
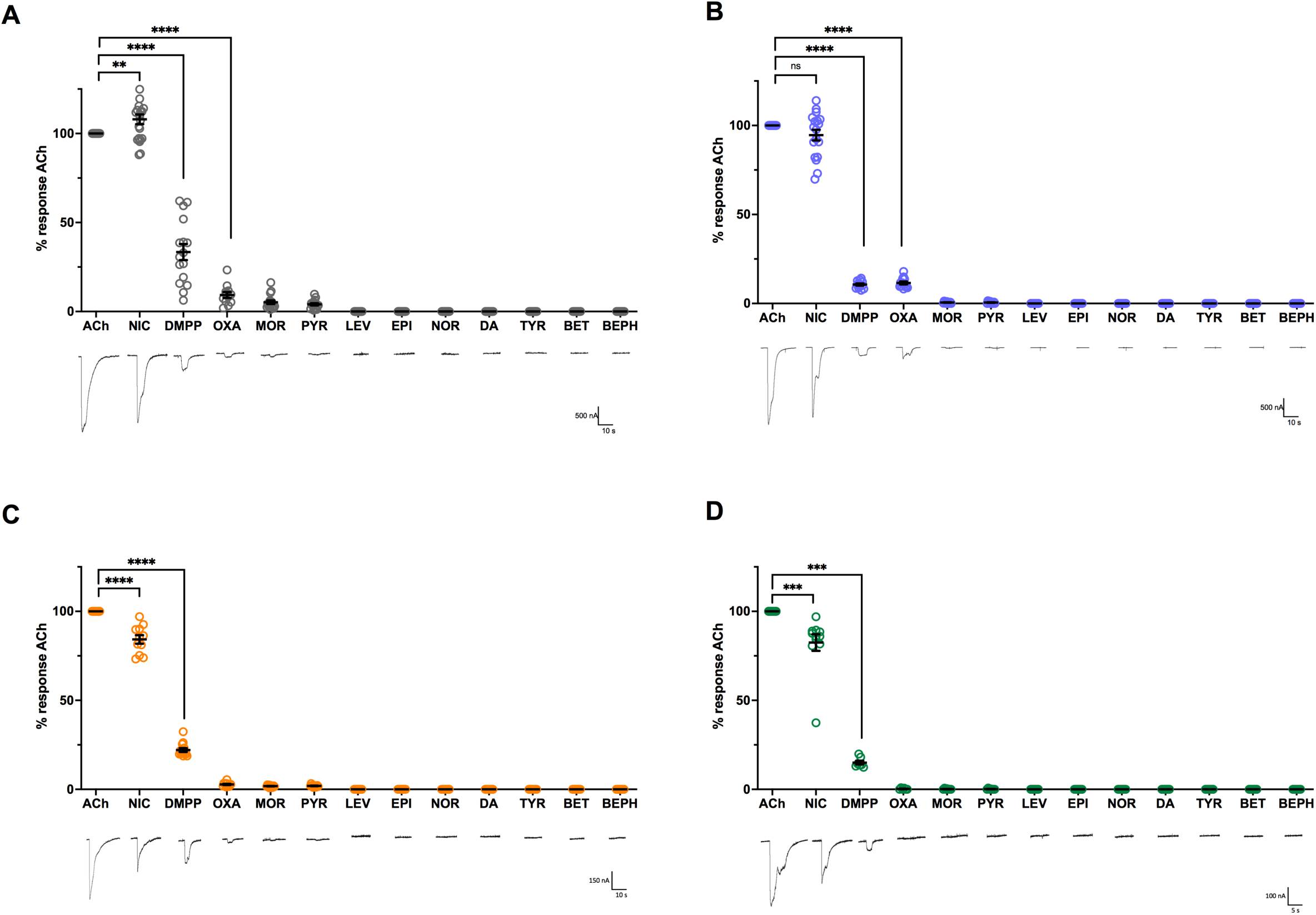
No change in ligand response profile is observed in clade III N-AChRs. N-AChR responses to different pLGIC agonists were measured to determine if declining maximal response is due to changing ligand preference. Current responses to 100 µM for all ligands are shown normalized to acetylcholine for (A) *A. suum*, (B) *D. medinensis*, (C) *G. pulchrum*, (D) *T. callipaeda*. All four receptors had similar response profiles: ACh ≥ NIC >> DMPP> OXA = MOR = PYR > LEV= EPI = NOR = DA = TYR = BET = BEPH. Circles represent responses from individual oocytes. Similar ligand response profile indicates no change in ligand occurs down clade III receptor phylogeny. **p<0.01; ***p<0.0005; ****p<0.0001. Error bars represent standard error. n>6 for all.

### Later clade III N-AChRs take longer to reach maximal response

Current responses were normally measured two days after oocyte injection. Any decreasing efficiency in receptor production in later clade III N-AChR would mean receptor synthesis, and therefore maximal measurable current, would occur at a later time point in more derived receptors. AChRs have different expression times depending on the receptor (typically 2 days for the N-AChR and 5 days for the L-AChR) [1,33,47,48]. To identify the time taken to reach maximal current, ACh-induced responses were measured each day after injection of receptor cRNA, for three consecutive days and normalized to responses obtained on Day 3. Receptors for each species took progressively longer to reach maximal response along the phylogeny (Fig 2C). After Day 1, both Asu-ACR-16 and Dme-ACR-16 had nearly half of their Day 3 maximal response; 44 ± 5%, p<0.0001 and 49 ± 7% p<0.0001, respectively. By Day 2 both receptors had reached responses that were no longer significantly smaller than Day 3, 88 ± 5%, p = 0.0841 for Asu-ACR-16 and 102 ± 7%, p=0.7011 for Dme-ACR-16. This in contrast to *G. pulchrum* and *T. callipaeda* that did not reach a maximum by Day 2. On Day 1, Gpu-ACR-16 had only 19 ± 6 %, p<0.0001 of its Day 3 response, and on Day 2 had reached 76 ± 6%, p<0.05 response. On Day 1, Tzc-ACR-16 had 7 ± 1% p<0.0005 of its Day 3 response, and by Day 2 had only reached 66 ± 21 % p<0.05. These results indicate that earlier clade III N-AChRs reach maximal response more efficiently compared to later clade III receptors.

### *B. malayi* ACR-16 requires at least three additional accessory proteins

No meaningful difference in the way the N-AChR from different species responded to various agonists, coupled with delayed maximal responses in more derived receptors suggested that a decrease in overall receptor function was likely responsible for the decreased response. The nicotine response of *B. malayi in vivo* [29,42], the failure to reconstitute a receptor *ex vivo*, together with a delayed time to maximal response for the species closest to *B. malayi* suggests a change that affected expression *ex vivo* specifically. This could be due to an increased dependence one or more members of the network of regulatory accessory proteins that were not included in the cRNA injected into oocytes. A review of accessory proteins known to be involved in *C. elegans* AChR expression identified UNC-50, UNC-74, MOLO-1, EAT-18, EMC-6, NRA-2 and NRA-4 as possible candidates [1,39,40,49,50]. These accessory proteins were then included with each N-AChR individually or in a series of combinations to determine if a specific combination was needed to produce the *B. malayi* N-AChR. UNC-50 and UNC-74 were co-injected because of their joint requirement for expression of the L-AChR in oocytes [1,48,51]. NRA-2 and NRA-4 were co-injected because they function in tandem for subunit assembly [40].

Receptor responses in the presence of different accessory protein conditions are shown in Fig 3C, S1 Fig and S1 Table. The response of each receptor with *B. malayi* RIC-3 was used as the reference condition for the other accessory protein conditions. In general, the same progressive decline was observed, regardless of accessory protein combination, however some combinations produced significant deviations from the *B. malayi* RIC-3 reference condition. Responses for Asu-ACR-16 were higher under all additional accessory protein conditions except for when MOLO-1 was added which decreased the response (1525 ± 5 nA, p<0.0001, S1 Fig). The absence of any accessory protein produced an inconsistent, little to no response (24 ± 10 nA, p<0.0001, S1 Fig). None of the additional accessory proteins affected Dme-ACR-16 responses. Interestingly, when no accessory protein was included, it still produced large consistent current (1290 ± 70 nA, p<0.0001, S1 Fig). Current responses for Gpu-ACR-16 were increased when RIC-3, UNC-50, UNC-74 and EAT-18 were combined (1854 ± 83, p<0.01, S1 Fig) and when RIC-3, NRA-2 and NRA-4 were combined (2553 ± 123 nA, p<0.0001, Fig 3C). As expected, current responses were completely inhibited without any RIC-3 (0 ± 0 nA, p<0.0001, S1 Fig). Current responses for Tzc-ACR-16 were decreased when RIC-3, UNC-50 and UNC-74 were combined (367 ± 43nA, p<0.0005, Fig 3C), increased when RIC-3, NRA-2 and NRA-4 were combined (1263 ± 88 nA, p<0.01, Fig 3C) and abolished when no accessory protein was included (0 ± 0 nA, p<0.0001, S1 Fig). BMA-ACR-16 only produced measurable currents when RIC-3 + EMC-6 (17 ± 4, p<0.01, Fig 3C) or RIC-3, NRA-2 + NRA-4 (4 ± 2, p<0.05, Fig 3C) were combined. Therefore, EMC-6, NRA-2, and NRA-4 accessory proteins play a more critical role for functional expression of more derived clade III receptors. This is seen by decreased receptor response in a heterologous system where the native accessory proteins are not present.

The increased dependence on EMC-6, NRA-2 and NRA-4 suggests an evolved change in the ACR-16 subunit itself. In order to determine approximately where in the subunit sequence these changes occur, a series of chimeras were made between *A. suum* and *T. callipaeda* ACR-16 by exchanging each of their three main structural regions (S2A Fig). *T. callipaeda* was chosen rather than *B. malayi* so that the small current observed for *T. callipaeda* could be used to verify that the chimeras were in fact functional. The response of the chimeras was not straightforward, however they suggested all main structural regions contribute to the reduced responses to varying degrees, with the ECD having the largest effect (S2C Fig). Replacing the *T. callipaeda* ECD into Asu-ACR-16 (TZC-ECD) inhibited nearly all response to ACh (42 ± 9 nA, p<0.0001) whereas presence of the *A. suum* ECD in Tzc-ACR-16 (ASU-ECD) significantly increased the response (1819 ± 115 nA, p<0.0001). Both ICL chimeras produced large responses. The *T. callipaeda* ICL in Asu-ACR-16 (TZC-ICL) was not significantly different from native Asu-ACR-16 (1959 ± 78 nA, p=0.593). The *A. suum* ICL in Tzc-ACR-16 (ASU-ICL) significantly increased the response (2237 ± 98 nA, p<0.0001). The *A. suum* TM regions in Tzc-ACR-16 (ASU-TM) severely decreased maximal response (24 ± 8 nA, p<0.0001), while the Tzc-TM region in the *A. suum* ACR-16 (TZC-TM) increased the response (2917 ± 105 nA, p<0.0001). To summarize, later clade III N-AChR current responses can be increased with the addition of new accessory proteins EMC-6 or NRA-2 + NRA-4, and they are sufficient to produce a measurable *B. malayi* N-AChR. Chimeras indicate that all regions may contribute to this requirement, with the ECD having a majority of the effect.

### Pharmacology of chimeric *B. malayi* ACR-16

The original goal of this work was to characterize the response of the *B. malayi* N-AChR. Initially no response could be seen from Bma-ACR-16. Addition of EMC-6, NRA-2 and/or NRA-4 led to a detectable response to ACh but the currents remained small. The highest responses for Bma-ACR-16 were obtained with RIC-3 + NRA-2 (33 ± 9 nA, p<0.05, Fig 6B), RIC-3 + NRA-4 (31 ±13 nA, p=0.0698, Fig 6B) or RIC-3 + EMC-6 (17 ± 4 nA, p<0.05, Fig 3C). Other combinations reduced measurable currents; RIC-3 + NRA-2 + NRA-4 + EMC-6 (11 ± 9 nA, n=11, p=0.3108, Fig 3C) and RIC-3 + NRA-2 + NRA-4 EMC-6 + EAT-18 (2 ± 1 nA, p=0.2019, Fig 6B), and one case abolished the response entirely RIC-3 + NRA-2 + NRA-4 + EMC-6 + EAT-18 + UNC-50 + UNC-74 (0 ± 0 nA, Fig 6B). A similar phenomenon has been seen previously for other N-AChRs [1,32,36].

**Fig 6.**
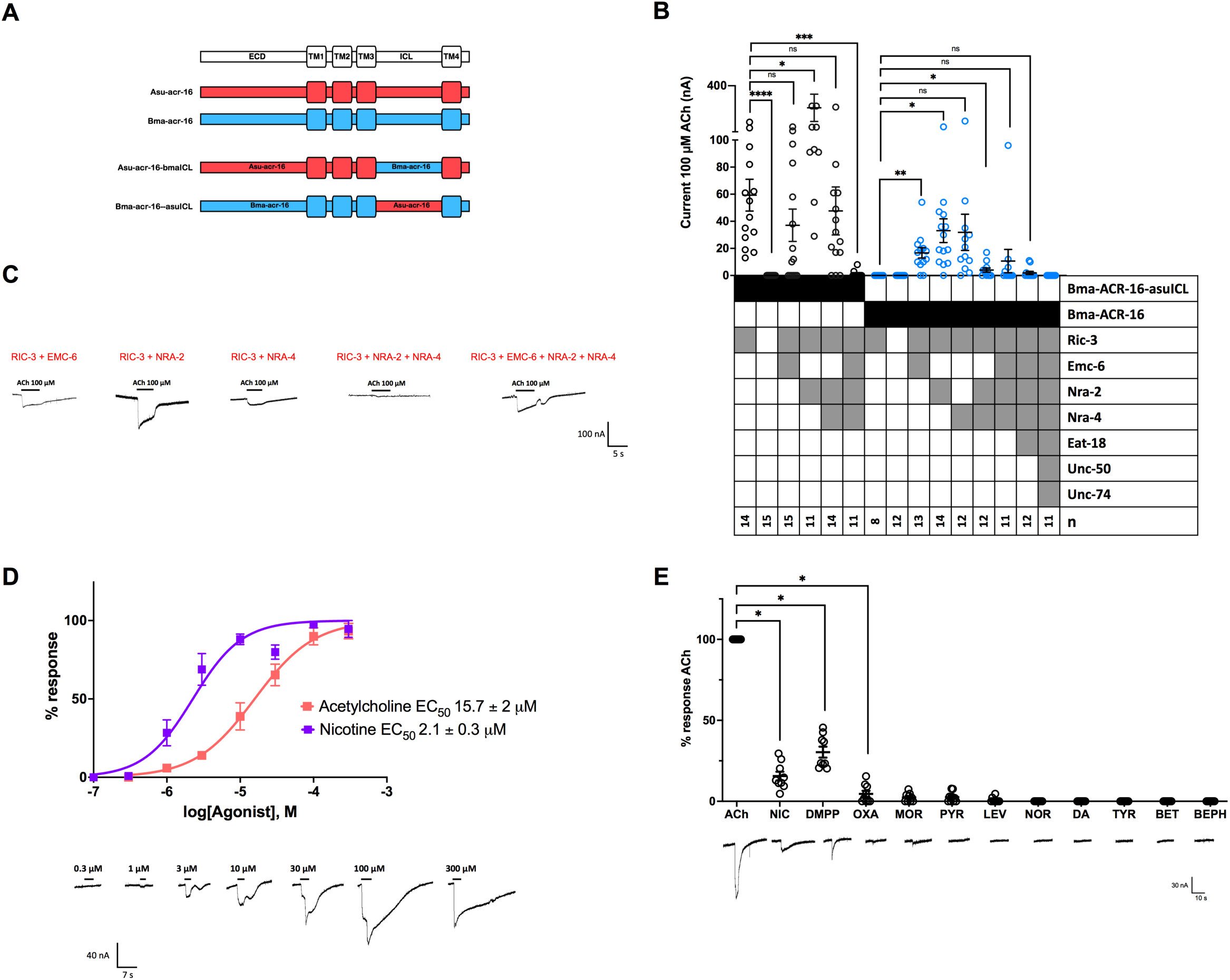
The *B. malayi* N-AChR requires additional accessory proteins and has unchanged pharmacology compared to other clade III N-AChR. (A) Bma-ACR-16-asuICL and Asu-ACR-16-bmaICL chimeras. (B) Bma-ACR-16-asuICL and Bma-ACR-16 were co-injected with a variety of accessory protein combinations to identify those that produces the highest current response. Bma-ACR-16-asuICL chimera in black, Bma-ACR-16 in blue. Circles represent measured currents from individual oocytes to 100 µM acetylcholine. Bma-ACR-16 responses (compared to RIC-3) to different combinations of EMC-6, NRA-2 and NRA-4 were: RIC-3 + NRA-2 (33 ± 9 nA, p<0.05); RIC-3 + NRA-4 (31 ±13 nA, p=0.0698); RIC-3 + EMC-6 (17 ± 4 nA, p<0.05) ; RIC-3 + NRA-2 + NRA-4 EMC-6 (11 ± 9 nA, n=11, p=0.3108); RIC-3 + NRA-2 + NRA-4 EMC-6 + EAT-18 (2 ± 1 nA, p=0.2019); and RIC-3 + NRA-2 + NRA-4 + EMC-6 + EAT-18 + UNC-50 + UNC-74 (0 ± 0 nA). Bma-ACR-16-asuICL chimera responses under the different accessory protein conditions were: RIC-3 (59 ±43 nA); RIC-3 + NRA-2 (256 ± 295 nA, p<0.05); RIC-3 + NRA-2 + NRA-4 (47 ± 66 nA, p=0.5887); RIC-3 + EMC-6 (37 ± 46 nA, p=0.1957); RIC-3 + NRA-2 + NRA-4 + EMC-6 (1 ± 2 nA, p<0.005). RIC-3 + NRA-2 was therefore included for acetylcholine and nicotine dose-response curves and for ligand panel analysis. Error bars represent standard error. *p<0.05; **p<0.01; ***p<0.0005; ****p<0.0001. (C) Sample recordings of Bma-ACR-16 with various accessory protein conditions in response to 100 µM acetylcholine. (D) Dose-response curves were measured for the Bma-ACR-16-asuICL chimera to compare to the other clade III ACR-16. Dose-response curves for acetylcholine in pink (EC_50_ = 15.7 ± 2 µM, n=6) and nicotine in purple (EC_50_ = 2.1 ± 0.3 µM, n=8). Hill coefficients were 1 ± 0.1 for ACh and 1.2 ± 0.1 for NIC. Sample recording to acetylcholine is shown. Similar EC_50_ to other clade III ACR-16 indicates the small currents obtained with Bma-ACR-16 is not due to changes in ligand affinity. Error bars represent standard error. (E) Bma-ACR-16-asuICL chimera response to 100 µM ligands relative to acetylcholine. Bma-ACR-16-asuICL chimera had a comparable response profile to other N-AChR. Relative responses to ACh were NIC (16 ± 3 % p<0.05), DMPP ( 30 ± 3 %, p<0.05), OXA (5 ± 2 %, p<0.05), MOR (2 ± 1 %, p<0.05), and PYR (3 ± 1 %, p<0.05), LEV (1 ± 1 %, p<0.05),) and no responses to NOR (0 ± 0 %, p<0.05), DA (0 ± 0 %, p<0.05), BET (0 ± 0 %, p<0.05), BEPH (0 ± 0 %, p<0.05) and TYR (0 ± 0 %, p<0.05). Sample recording to each ligand is shown. Similar ligand response profile to other clade III N-AChRs indicates the small currents obtained with Bma-ACR-16 is not due to changes in ligand preference. Error bars represent standard error. *p<0.05.

The ability of the *A. suum* ACR-16 ICL to increase overall response in the reciprocal chimera (S2C Fig) suggested that a chimera of Bma-ACR-16 containing the *A. suum* ICL (Bma-ACR-16-asuICL) may produce larger responses overall to be used for *B. malayi* N-AChR pharmacology characterization since ligand-interaction and gating are determined predominantly by the ECD and TM regions [9,52–55]. Chimeras where the ICL has been exchanged have previously shown not to affect receptor pharmacology [56,57]. A comparison between the Asu-ACR-16 and Asu-ACR-16 containing the *B. malayi* ICL (Asu-ACR-16-bmaICL) was used as a control to verify that the receptor response was in fact determined by the ECD and TM regions (S3 Fig). Asu-ACR-16-bmaICL EC_50_ for ACh was 4.3 ± 0.5 μM and for NIC was 4.3 ± 0.5 μM (S3C Fig) and had similar ligand response profile as the full-length Asu-ACR-16 receptor (S3D Fig). The Asu-ACR-16-bmaICL chimera’s similarity in pharmacology to the native Asu-ACR-16 receptor confirms that the ICL exchange in the Bma-ACR-16 chimera provides accurate receptor pharmacology.

The Bma-ACR-16-asuICL chimera responses were small, but RIC-3 + NRA-2 produced the largest and most reproducible currents (256 ± 295 nA, p<0.05, Fig 6B). Other combinations were smaller; RIC-3 + NRA-2 + NRA-4 (47 ± 66 nA, p=0.5887, Fig 6B); and RIC-3 + EMC-6 (37 ± 46 nA, p=0.1957, Fig 6B), and RIC-3 + NRA-2 + NRA-4 + EMC-6 (1 ± 2 nA, p<0.005, Fig 6B) (see Fig 6C for sample recordings). Receptor characterization was therefore carried out with the Bma-ACR-16-asuICL chimera co-injected with RIC-3 + NRA-2 and its pharmacology was similar to other N-AChR studied here. The EC_50_ for ACh and NIC of the Bma-ACR-16-asuICL chimera was 15.7 ± 2 μM and 2.1 ± 0.3 μM respectively (Fig 6C). Relative responses to ACh (Fig 6D) were NIC (16 ± 3 % p<0.05), DMPP ( 30 ± 3 %, p<0.05), OXA (5 ± 2 %, p<0.05), MOR (2 ± 1 %, p<0.05), and PYR (3 ± 1 %, p<0.05), LEV (1 ± 1 %, p<0.05),) and no responses to NOR (0 ± 0 %, p<0.05), DA (0 ± 0 %, p<0.05), BET (0 ± 0 %, p<0.05), BEPH (0 ± 0 %, p<0.05) and TYR (0 ± 0 %, p<0.05). To summarize, we were successful in measuring the first recombinant *B. malayi* AChR by including new accessory proteins EMC-6, NRA-2 and NRA-4. Furthermore, our characterization of a chimeric Bma-ACR-16 confirmed this acquired accessory protein requirement occurs independent of pharmacology. Despite the success in measuring the *B. malayi* N-AChR, its small current responses suggest other accessory proteins are still missing.

## Discussion

Reconstitution of anthelmintic target receptors *ex vivo* has proven critical for understanding the neuromusculature of nematodes. While the response of tissues [30,49], and even single receptors can be determined *in vivo* [46,58,59], it remains challenging to identify the specific components of these receptors and their detailed characteristics [1]. The N-AChR, a homopentamer of ACR-16 has been characterized through reconstitution for many different nematodes, both parasitic [32–34,36,37] and free-living [24]. An initial attempt to characterize the *B. malayi* N-AChR failed when co-expressed with RIC-3, an ER localized accessory protein involved in subunit assembly required for other species N-AChR [1,25,32–36,60]. The absence of stop codons or amino acid substitutions at positions known to be critical for receptor function suggested the failure may not have been an experimental artefact.

*A. suum* produces a robust N-AChR current in *Xenopus* oocytes [36] and is basal to the clade of filarial nematodes to which *B. malayi* belongs [38]. We therefore characterized five clade III N-AChRs: *A. suum*, *B. malayi*, and three intermediate species in order of phylogeny, *D. medinensis, G. pulchrum* and *T. callipaeda*. These species represent independent samples of the ancestral state between *A. suum* and *B. malayi* ACR-16 and so may shed light on the changes leading to an inability to reconstitute the receptor. Two possibilities were expected, either a discrete change that would suggest a single mutational event may be responsible, or a gradual decline that would suggest an adaptive change. The maximal response to ACh and NIC declined progressively through the clade of filarial nematodes, with Asu- and Dme-ACR-16 responding the most (∼2000 - 2200 nA), followed by Gpu-ACR-16 (∼1400 nA), then Tzc-ACR-16 (∼400 - 600 nA) and Bma-ACR-16 with no measurable current. Therefore, the change appears to be an evolved characteristic.

The progressive decrease in measurable response could be due to a decreased affinity for the activating ligands, possibly with changes in the profile of response to other agonists. This could be ruled out for all since the EC_50_ for ACh and NIC did not change meaningfully between species and were similar to the other N-AChRs characterized so far (Table 1) [36]. In addition, the relative order of response to a panel of agonists was the same for each species. This is consistent with the fact that a response to nicotine in living *B. malayi* is found and its characteristics are unremarkable [29,42].

**Table 1.** Comparison of pharmacology and accessory proteins required for nematode N-AChR.

| | | Approximate response | EC50 μM ACh | EC50 μM NIC | Accessory protein | Source |
| --- | --- | --- | --- | --- | --- | --- |
| Clade II | *T. suis* | 7000 nA | $14.5 \pm 1.03$ | - | Ric-3 | [34] |
| | *T. muris* | 10,000 nA | $31.1 \pm 0.3$ | - | Ric-3 | [35] |
| Clade V | *C. elegans* | 7000 nA | $22.0 \pm 1.2$ | $15.1 \pm 1.3$ | Ric-3 | [1,24,37]<sup>a</sup> |
| | *N. americanus* | 1000 nA | $170.1 \pm 19.23$ | $597.9 \pm 59.12$ | Ric-3 | [33] |
| | *A. celanicum* | 2000 nA | $20.64 \pm 0.32$ | $24.37 \pm 2.89$ | Ric-3 | [33] |
| | *A. caninum* | 100 nA | $4.3 \pm 0.0$ | - | Ric-3 | [32] |
|  | *H. contortus* | No response | - | - | - | [37] |
| Clade III | *P. equorum* | 7000 nA | $6.4 \pm 0.6$ | $2.9 \pm 0.5$ | Ric-3 | [37] |
| | *A. suum* | 2000 nA | $8.9 \pm 0.3$ | $5.1 \pm 0.2$ | Ric-3 | [36] Present study<sup>b</sup> |
| | *D. medinensis* | 2000 nA | $12.8 \pm 0.4$ | $3.8 \pm 0.2$ | none | Present study |
| | *G. pulchrum* | 1000 nA | $10.5 \pm 0.5$ | $2.1 \pm 0.1$ | Ric-3 | Present study |
| | *T. callipaeda* | 700 nA | $17.4 \pm 0.8$ | $4.9 \pm 0.4$ | Ric-3 | Present study |
| | *B. malayi chimera* | 20-200 nA | $15.7 \pm 2$ | $2.1 \pm 0.3$ | Ric-3, emc-6, nra-2, nra-4 + others? | Present study<sup>c</sup> |

All nematode N-AChR reconstituted *ex vivo* require the co-injection of RIC-3 except for *B. malayi* that requires the addition of at least 3 others and *D. medinensis* that does not require any. Similar pharmacological profile of clade III receptors, coupled with smaller responses and additional accessory protein requirement in later clade III N-AChR, suggests that progressive accessory protein differences occur independently from pharmacology.
<sup>a</sup> values shown for *C. elegans* are from Charvet et al., 2018 [37].
<sup>b</sup> values shown for *A. suum* are from this study, not the original Asu-ACR-16 characterization [36].
<sup>c</sup> values are shown for the *B. malayi* ACR-16 chimera.

An alternative to pharmacology changes could be that fewer receptors were expressed at the surface of oocytes *ex vivo*. If so, the ACR-16 subunit may have become more dependent on components of the processing and transport machinery in *B. malayi* to the point where receptor assembly and surface expression cannot be measured without additional components *ex vivo*. This is true more generally for the L-AChR that requires three accessory proteins co-injected within the oocyte system to produce measurable currents [1,14,48,61,62]. An increased dependence on missing factors would make expression less efficient and so take longer for receptors be expressed at the surface. Our expression analysis further supported this idea with progressively longer times to reach maximal responses. Asu- and Dme-ACR-16 reached maximal currents by Day 2 whereas Gpu- and Tzc-ACR-16 took longer.

Characterization of accessory proteins associated with expression of cationic pLGICs in nematodes has focused primarily on the L-AChR of *C. elegans*. However, since the N-AChR and the L-AChR are expressed in the same tissue in *C. elegans* [3] and other nematodes including *B. malayi* [29,31], we hypothesized that ACR-16 would potentially interact with those that are co-expressed *in vivo*. A literature search for characterized *C. elegans* L-AChR accessory proteins identified seven: UNC-50, UNC-74, MOLO-1, EAT-18, EMC-6, NRA-2 and NRA-4 [1,39,40,49,50,63]. We also included EAT-18 that is expressed in the *C. elegans* pharynx and is required for expression of the EAT-2 receptor that is within the ACR-16 clade [50,64]. *B. malayi* has lost the gene encoding *eat-2* [41] and retention of *eat-18* may suggest it has acquired other functions. Only the inclusion of EMC-6 or NRA-2 + NRA-4 together with RIC-3 resulted in a measurable *B. malayi* N-AChR.

Our strategy to explore the reason for failure to reconstitute a *B. malayi* N-AChR was to examine intermediate species between *A. suum,* which reconstitutes well, and *B. malayi*. The progressive decreased response towards *B. malayi* suggested that the phenomenon was one extreme of an adaptive response within the filarial nematodes, rather than a discrete event limited to *B. malayi*. This appears confirmed by the fact that inclusion of NRA-2 and NRA-4 led to a detectable Bma-ACR-16 response and also increased the response of Gpu-ACR-16 and Tzc-ACR-16. Inclusion of EMC-6 also produced a Bma-ACR-16 response. The largest response from the *B. malayi* N-AChR occurred in combinations of RIC-3 with NRA-2 or NRA-4. Even so, the response was far below what can be expected for an N-AChR expressed in oocytes. We expect other, as yet unidentified accessory proteins are required to obtain a robust receptor *ex vivo*.

Addition of other accessory proteins that were tried here reduced the observed response. A similar phenomenon has been reported previously for other N-AChRs [1,32,36]. We know that the receptor synthesis machinery is complex and involves a network of more than 200 proteins [23]. Interaction between these proteins may influence the overall expression of a specific channel. The presence of only a few of these in the oocytes may dis-regulate their normal function and lead to decreased expression. Two species-specific effects of accessory proteins were also observed. First, MOLO-1 significantly decreased responses for Asu-ACR-16, but not the others. This suggests MOLO-1 interacts with the Asu-ACR-16 receptor in such a way that may decrease the open probability of the channel at the surface membrane [49]. Second, *D. medinensis* N-AChR no longer required any RIC-3 for expression in oocytes. This was unexpected because all previously characterized N-AChRs require RIC-3 [1,32–37] and indicates that specific changes occurred in Dme-ACR-16 altering its interaction with RIC-3.

The increased dependence on accessory proteins for the N-AChR in the filarial nematodes is novel. Since the accessory proteins included were the same for each species, the change in requirement must be due to the specific subunit sequence. Chimeras are a common tool for identifying functional regions within receptor subunits [55,65,66]. Associating the receptor phenotype with the presence of a specific region can infer that the relevant sequence lies within that region. In this case, the phenotype associated with the *B. malayi* N-AChR is an absence of response that would be indistinguishable from a cloning artefact that led to a non-functional construct. For this reason, the major subunit structural domains were exchanged between *A. suum* and *T. callipaeda* ACR-16. These chimeras suggested all regions may contribute to the change in maximal responses. Exchanging the ECD alone reversed the receptor response indicating that the ECD confers the majority of the increased accessory protein requirement. This coincides with the fact that many expression and synthesis signals are found within this region [67–69]. Exchanging the ICL alone was not clear since the constructs with either the Asu-ICL or Tzc-ICL both produced a relatively high response. This may be an indication that a second region conferring increased dependence on additional accessory proteins exists within the ICL, but that it requires a matching region. This may be due to the region being distributed to some extent rather than being discrete. Regardless, the eukaryotic ICLs are highly variable in sequence [11], and their precise structure and function remains unclear. The chimera results presented here agree with previous work showing that the ICL may be involved in regulating receptor synthesis and/or interactions with accessory proteins [70–72].

We could expect that an increased dependence for specific accessory proteins may be due to a physical interaction with them, and our chimeras identified that the majority of the decline is located in the ECD. Interestingly, few amino acid changes were found within the ACR-16 ECD. The Bma-ACR-16 ECD has 9 amino acids substitutions with the Tzc-ACR-16 ECD and 16 with the Asu-ACR-16 ECD. This implies few amino acid changes have a large effect on receptor synthesis and that some of these residues may be critical for receptor synthesis. EMC-6, NRA-2 and NRA-4 all act within various stages of ER receptor synthesis. EMC-6 is one protein within a larger ER membrane protein complex (termed EMC) that is conserved throughout the eukaryotes and is estimated to contain upwards of 10 different proteins [73]. The EMC functions as an insertase by directing the appropriate insertion of transmembrane domains through the membrane [74,75]. In *C. elegans*, EMC-6 knock-out led to decreased responses from all three body-wall muscle receptor types (L-AChR, N-AChR and UNC-49) and decreased expression of at least the L-AChR [39]. Including EMC-6 in oocytes may promote the appropriate folding and processing of the *B. malayi* N-AChR, thus increasing the amount of functional protein at the membrane surface. In its absence is it possible that the *B. malayi* N-AChR is not inserted into the membrane properly, which would target it for degradation. Five other EMC-6 paralogs are predicted in the *C. elegans* genome [39], and it is possible their co-injection may further increase *B. malayi* N-AChR responses in oocytes. NRA-2 is a nicalin-like homolog and NRA-4 is a NOMO-like homolog [23,40]. In *B. malayi*, NRA-2 is thought to sense intracellular influx of calcium and regulate the levels of the different L-AChR subunits to produce receptors with different subunit combinations [42]. In *C. elegans*, NRA-2 and NRA-4 KO worms had reduced L-and N-AChR responses and the L-AChR had altered properties [40]. It was proposed that they function together by determining the subunits that form a channel and exit the ER [40]. In another study, NRA-2 was found to act as a quality control check to retain mutant DEG/ENaC channels in the ER [76]. Their precise function remains unclear but based on current knowledge, including NRA-2 and NRA-4 with more derived clade III N-AChR may promote correct assembly and/or exit of receptors in the ER.

The ultimate goal of characterizing the *B. malayi* N-AChR remained challenging due to the small responses observed (<35 nA). To overcome this a chimera of Bma-ACR-16 containing the *A. suum* ICL (Bma-ACR-16-asuICL) was characterized instead when co-expressed with RIC-3 and NRA-2. We were able to confirm that the complementary chimera of Asu-ACR-16 containing the *B. malayi* ICL was not substantially different from the Asu-ACR-16, as expected from other similar studies [9,52–54]. These chimeras therefore provide accurate estimates of receptor pharmacology. The Bma-ACR-16-asuICL chimera had similar EC_50_ and agonist response profiles compared to other clade III receptors suggesting that *B. malayi* ACR-16 indeed functions as a homomeric nicotinic acetylcholine receptor with an increased dependence on accessory proteins.

Although an increased dependence on additional accessory proteins appears to be a feature of N-AChRs within the filarial nematodes, the reason for this remains unclear. One possibility is that it is a response to the many pLGIC subunit gene loss and duplication events observed within this clade. The filarial parasites have lost subunit genes *acr-2*, *acr-3* and *lev*-*1* and have duplicated *unc*-38 and *unc*-*63* [7,38,77]. Subunits that comprise pLGICs are subject to a rigorous synthesis pathway with as little as 5 - 30% of subunits becoming mature receptors at the cell surface [78,79]. A changing receptor subunit landscape means changes to how the subunits interact with each other, potentially leading to increased production of non-functional receptors. Structural change in subunit interaction may be enabled through accessory proteins that stabilize assembly intermediates. This dependence may have led to subunits that enhance the interaction with accessory proteins that would manifest as an increased requirement *ex vivo*. This could suggest a dynamic and inter-dependent relationship between receptor subunits and their synthesis machinery. This agrees with the fact that all three additionally required accessory proteins act within the subunit folding and/or assembly stages of expression.

Here we present a novel phylogenetic approach to studying receptor change. By characterizing N-AChRs from intermediate species, we were able to show a progressive increase in dependence for EMC-6, NRA-2 and NRA-4 in filarial worms, determined most predominantly by the ECD. Using this approach we have successfully measured the first reconstituted *B. malayi* AChR while also characterizing three new N-AChRs. The N-AChR characteristics of filarial nematodes has not changed substantially along with this change in accessory protein requirement. Focusing future efforts on proteins related to EMC-6, NRA-2 and NRA-4 may uncover other accessory proteins that increase overall expression of the N-AChR and possibly other AChRs from *B. malayi ex vivo*.

This work highlights the importance of accessory proteins for this receptor family in a heterologous system. Our comparative approach, together with the extensive genome data available for the nematodes, provides an approach that may prove fruitful to investigate the basis for other functional differences in neuromuscular drug target receptors.

## Experimental Procedures

### ACR-16 alignment and phylogeny

*Acr-16* sequences of the pLGIC family were identified from WormBase ParaSite WBPS13 [80,81]. The complete sequence was identified for each species except for *G. pulchrum*. Due to the poorer quality of its genome, eight amino acids (SGEWALPM) in the extracellular domain were missing from a region in the ECD that is identical in the other species and was inserted manually. ACR-16 subunit protein sequences were aligned in Geneious (v 9.0.5, Biomatters Ltd) using the MAFFT plugin (v7.017) [82]. The highly variable signal peptide and intracellular loop sections were removed. A maximum likelihood phylogeny was reconstructed using PhyML [83] with branch support values estimated from 1000 bootstrap resamples using the TBE option [84] and the BEST search option, with five random starting trees. A separate alignment of the ACR-16 studied here is shown in Fig 2.

Accession numbers of sequences used: Tca-acr-16 TCNE_0001320001-mRNA-1; Peq-acr-16 MH806894; Asu-acr-16 GS_23665 + GS_03796; Dme-acr-16 DME_0000017801-mRNA-1; Gpu-acr-16 annotated by hand; Tzc-acr-16 TCLT_0000979901-mRNA-1; Bma-acr-16 Bm13586a.1; Eel-acr-16 EEL_0000848201-mRNA-1; Lsi-acr-16 nLs.2.1.2.t06045-RA; Llo-acr-16 EJD73698.1; Avi-acr-16 nAv.1.0.1.t08089-RA; Ovo-acr-16 OVOC12002.1; Pex-acr-16 PEXSPEC000001098.t1; Ppa-acr-16 PPA34546.1; Hba-acr-16 Hba_18455 + Hba_18456 and annotated by hand; Hco-acr-16 HCON_00147330-00001; Nbr-acr-16 NBR_0001473901-mRNA-1 and annotated by hand; Dvi-acr-16 DICVIV_02269 and annotated by hand; Acn-acr-16 ACAC_0001143301-mRNA-1 and annotated by hand; Hbk-acr-16 HPOL_0000512601-mRNA-1; Ace-acr-16 Acey_s0049.g1864.t1; Aca-acr-16 ANCCAN_01899; Cjp-acr-16 CJA08620a.1; Cel-acr-16 F25G6.3.1; Cbr-acr-16 CBG01491.1

### Subunit and accessory protein cloning and RNA synthesis

*B. malayi acr-16* and accessory proteins *ric-3, unc-50, unc-74* and *molo-1* were cloned from adult female *B. malayi* cDNA received as a gift from Dr. T. Geary using the primers described in S1 Table. *B. malayi eat-18* was prepared from two overlapping oligonucleotide primers. Other subunits were synthesized from their predicted genome sequence (Gene Universal, USA) (S2 Table). *Asu-acr-16* and *Tzc-acr-16* sequences were re-engineered to provide unique restriction sites to generate chimeras that did not change the protein sequence (termed Asu-ACR-16-REM and Tzc-ACR-16-REM, respectively). All sequences were cloned into the pTD2 oocyte expression vector. Asu-ACR-16 and Tzc-ACR-16 chimeras were constructed using appropriate restriction enzymes, reciprocal exchange through ligation and confirmed by sequencing (McLab, USA).

cRNA for oocyte injection was prepared with the mMESSAGE mMACHINE T7 kit following manufacturers protocol (Ambion, USA). Concentration was determined using a Nanodrop spectrophotometer. Desired subunit combinations with associated accessory proteins were mixed and diluted with nuclease-free water to a final concentration of 250 ng/uL for each construct.

### Oocyte extraction and injection

Adult female *X. laevis* oocyte extraction was performed in accordance with the McGill University Animal Use Protocol 2015-7758 following standard protocol [85]. Briefly, frog ovary lobes were placed in a Ca^2+^-free OR2 solution and manually divided into clusters of <10 oocytes with thin tweezers followed by a 90-minute incubation in 10 mg/mL collagenase type II (Sigma-Aldrich, USA). Oocytes were washed in Ca2+-free OR2 and placed ND96 solution (NaCl 96 mM, KCl 2 mM, CaCl_2_ 1.8 mM, MgCl_2_ 1 mM and HEPES 5 mM, pH 7.3) supplemented with sodium pyruvate 2.5 mM [86] and incubated at 18°C until cRNA injection. Oocytes were injected with 50 nL (12.5 ng of each injected gene) of the desired cRNA subunit and accessory protein combination mix with the NanojectII (Drummond Scientific, USA). Following injection, oocytes were incubated in ND96 buffer at 18°C until electrophysiology. All electrophysiology was conducted two days post-injection, except for the expression analysis assay in which oocytes were measured every day after injection for three consecutive days.

### Pharmacological compounds

Receptor ligands acetylcholine (ACh), (-)-tetramisole hydrochloride (LEV), (-)-nicotine hydrogen tartrate (NIC), bephenium (BEPH), betaine (BET), norepinephrine (NOR), epinephrine (EPI), oxantel (OXA), morantel (MOR), pyrantel (PYR), tyramine (TYR), dimethylphenylpiperazinium (DMPP) and dopamine (DA) used for electrophysiology characterization were obtained from Sigma-Aldrich. All drugs were dissolved in recording solution (NaCl 100 mM, KCl 2.5 mM, CaCl2 1 mM, HEPES 5 mM, pH 7.3) to a stock concentration of 10 mM and diluted to their final recording concentrations. Oxantel and bephenium were first dissolved in DMSO to ensure a final recording concentration contained less than 0.1% DMSO. For agonist profiling, all drugs were diluted to a concentration of 100 μM. For EC_50_, NIC and ACh were diluted to concentrations between 0.3 and 300 μM. For maximum response measurements, 100 μM NIC and ACh was used.

### Electrophysiology

Oocytes were placed individually in an RC3Z chamber (Harvard Apparatus, USA) submerged in recording solution (NaCl 100 mM, KCl 2.5 mM, CaCl_2_ 1 mM, HEPES 5 mM, pH 7.3). The RC3Z chamber was connected to a perfusion system. Pulled glass capillaries (World Precision Instruments, USA) were filled with 3M KCl and the tips clipped and checked for appropriate resistance (between 0.5 and 5 MΩ). A 3M KCl agar bridge grounded the oocyte bath chamber. Measurements were made using an Axon Axoclamp 900A amplifier with Digidata1550M. Oocytes were voltage-clamped at -60 mV while exposed to drug solutions. Oocytes with holding potentials less than -400 nA were not considered for analysis due to poor membrane integrity. Data was analyzed using Clampex 9.2 (Axon Instruments, USA), statistical analysis and graphs produced in Prism (GraphPad version 9.1). T-tests were used to compare responses with error bars representing standard error.

## Acknowledgements

We thank Dr. Thomas Duguet for cloning *B. malayi acr-16* and Dr. Tim Geary for generously providing us with *B. malayi* cDNA.

## Supplemental Figure Legends

**S1 Fig.**
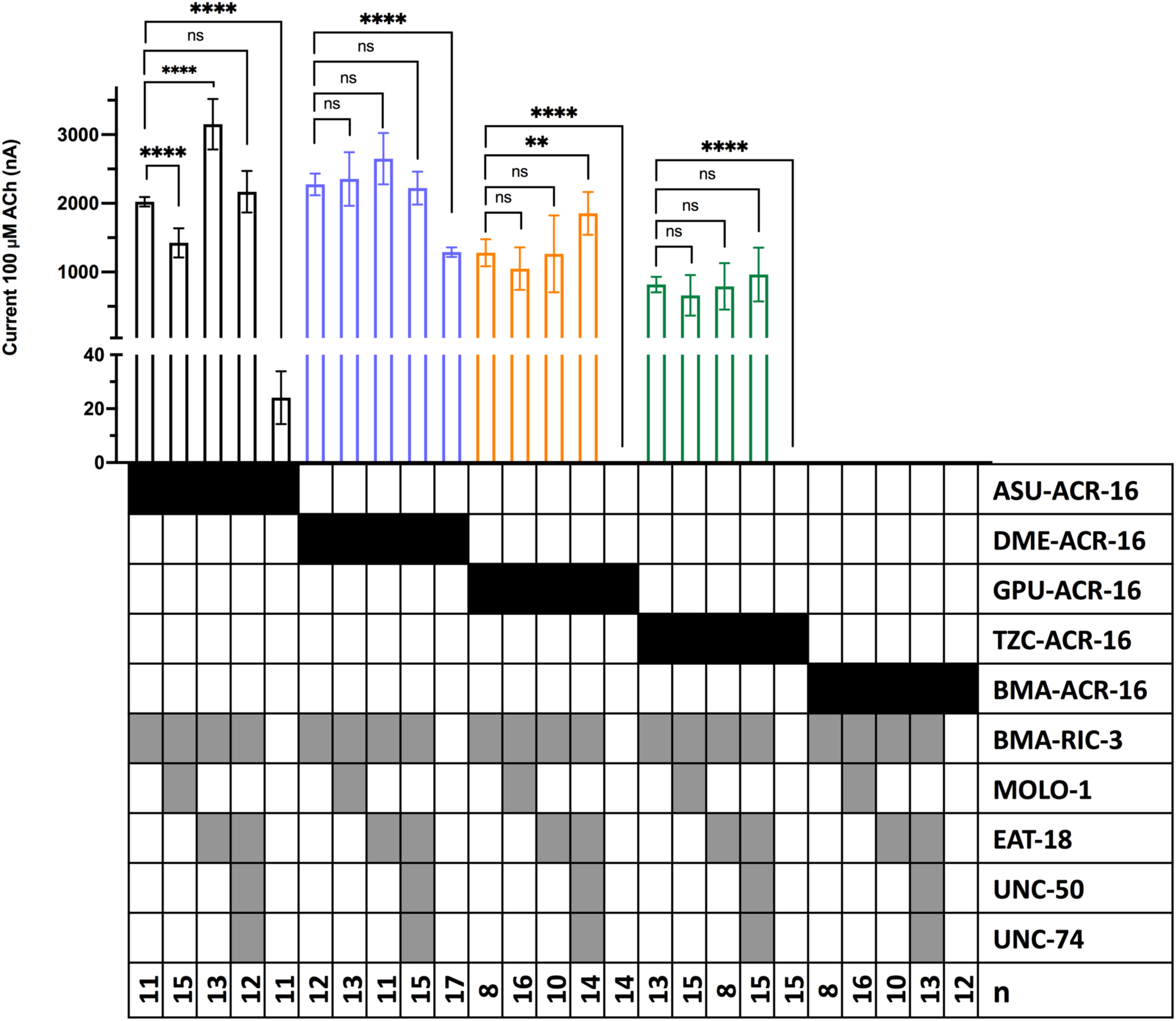
N-AChR responses with additional accessory protein combinations. Current responses of clade III N-AChRs *A. suum* (black), *D. medinensis* (purple), *G. pulchrum* (orange), *T. callipaeda* (green) and *B. malayi* (grey) to 100 µM ACh with different accessory proteins. Receptor responses are compared to Bma-RIC-3 condition. Error bars represent standard error. n>8, **p<0.01; ****p<0.0001. See S1 Table for values.

**S2 Fig.**
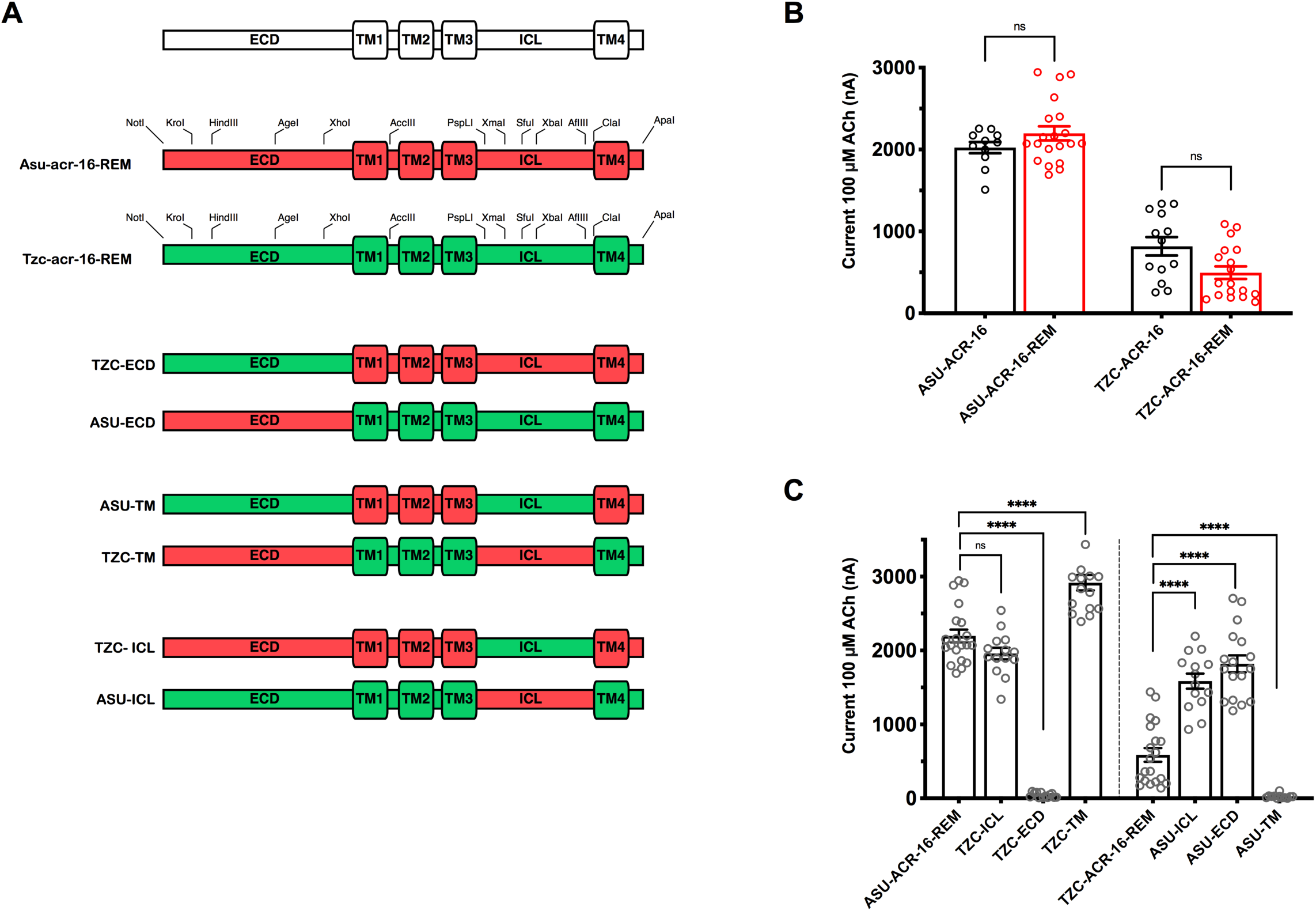
*A. suum* and *T. callipaeda* ACR-16 chimeras. The three main structural regions (ECD, ICL & TM) were exchanged between *A. suum* and *T. callipaeda* ACR-16 to identify the regions mediating the declining responses in later clade III N-AChRs. (A) *A. suum* and *T. callipaeda* ACR-16 coding sequences were modified to introduce restriction enzyme sites to constrict chimeras. *A. suum* shown in red and *T. callipaeda* in green. The modified sequences introduced complementary restriction enzyme sites termed ACR-16-REM. The three main structural regions (ECD, ICL, TMs) were exchanged between the two receptors. TZC-ECD contains the *A. suum* sequence with the *T. callipaeda* ECD. ASU-ECD contains the *T. callipaeda* sequence with the *A. suum* ECD. ASU-TM contains the *T. callipaeda* sequence with the *A. suum* TM. TZC-TM contains the *A. suum* sequence with the *T. callipaeda* TM. TZC-ICL contains the *A. suum* sequence with the *T. callipaeda* ICL. ASU-ICL contains the *T. callipaeda* sequence with the *A. suum* ICL. (B) Coding-sequence modified ACR-16 receptors (ACR-16-REM, red) produced responses indistinguishable from native ACR-16 sequence (black). Asu-ACR-16 (2023 ± 68 nA, n=11) versus Asu-ACR-16-REM (2197 ± 383 nA, n=20); and Tzc-ACR-16 (818 ± 112 nA, n=13) versus Tzc-ACR-16-REM (587 ± 415 nA, n=20). *B. malayi ric-3* co-injected with all. Circles represent currents measured from individual oocytes. Error bars represents standard error. (C) Chimeras exchanging each of the three structural regions between the receptors identified all regions contribute to the declining current responses measured in later clade III receptors. Circles represent currents measured from individual oocytes. Error bars represents standard error. Significance determined by comparing the current responses to the relevant reference receptor. ****p<0.0001.

**S3 Fig.**
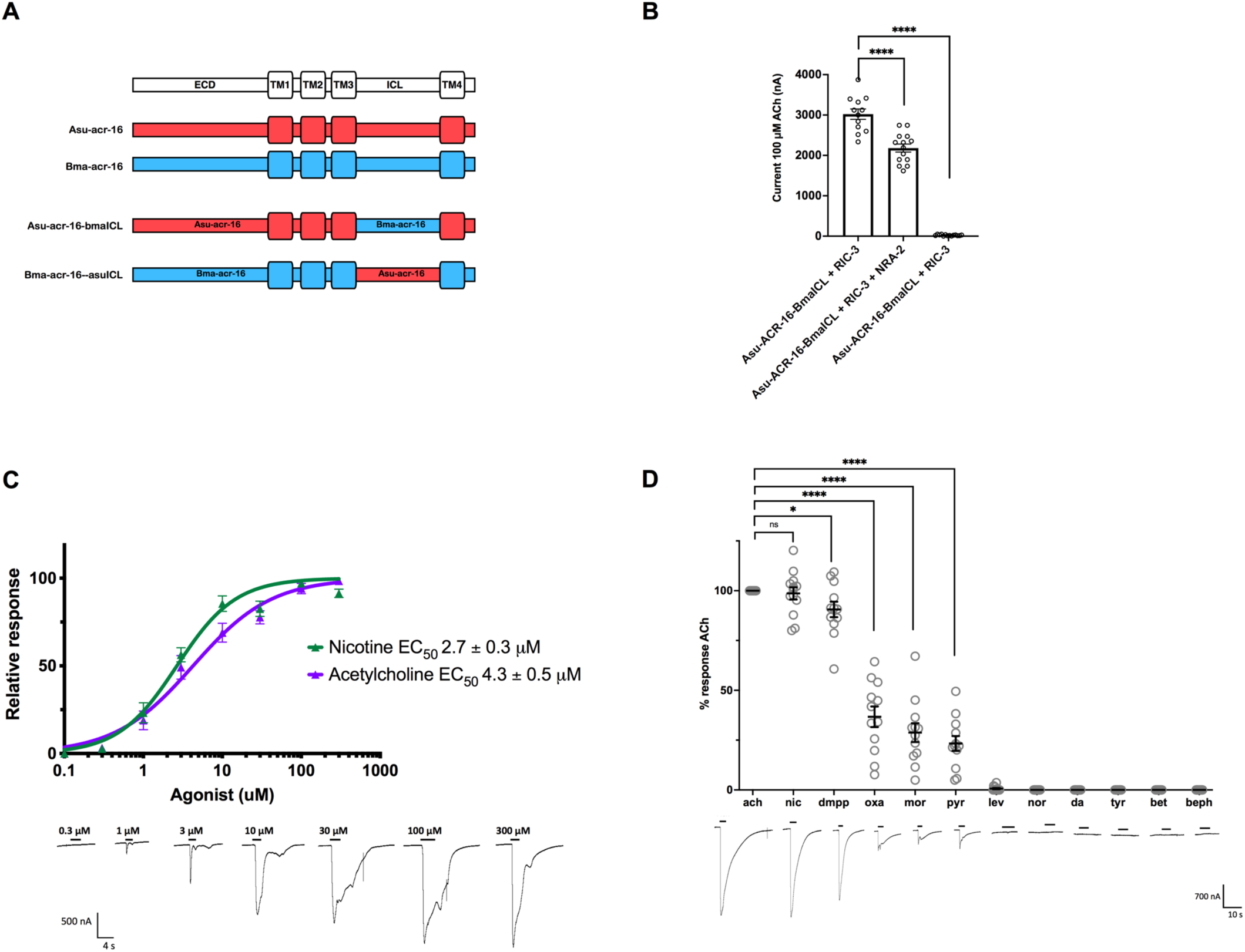
Asu-ACR-16-bmaICL chimera responses and pharmacology. (A) Chimeras for the *A. suum (red)* and *B. malayi acr-16 (blue) sequences.* Asu-ACR-16-bmaICL chimera has the *Asu-acr-16* subunit with the intracellular loop of *Bma-acr-16*, and Bma-ACR-16-asuICL chimera has the *Bma-acr-16* subunit with the intracellular loop of *Asu-acr-16*. Chimeras were made between *A. suum* and *B. malayi* ACR-16 to characterize *B. malayi* ACR-16 in the event that the unmodified *B. malayi* ACR-16 did not produce robust enough responses for characterization. The intracellular loop was exchanged since it is one of the regions thought to contribute to the declining responses observed in clade III ACR-16 and it does not contain the ligand-binding regions. (B) Asu-ACR-16-bmaICL chimera current responses under different accessory protein conditions: RIC-3 (3021 ± 444 nA, n=12); RIC-3 + NRA-2 (2181 ± 369 nA, n=14, p<0.0001); and no accessory protein (22 ± 15 nA, n=14, p<0.0001). Circles represent current responses from individual oocytes. Error bars represent standard error. ****p<0.0001. (C) Dose-response curves were measured for the Asu-ACR-16-bmaICL chimera to compare to the Asu-ACR-16 receptor as a control. Dose-response curves for acetylcholine in purple (EC_50_ = 4.3 ± 0.5 µM, n=8) and nicotine in green (EC_50_ = 2.7 ± 0.3 µM, n=10). Hill coefficients were 0.8 ± 0.1 for ACh and 1.2 ± 0.1 for NIC. Sample recordings to acetylcholine is shown. Error bars represent standard error. (D) Asu-ACR-16-bmaICL chimera response to ligands relative to acetylcholine. Concentrations of 100 µM was used for all ligands. Apart from higher responses to DMPP, Asu-ACR-16-bmaICL chimera had a comparable response profile compared to Asu-ACR-16: ACh = DMPP> NIC> OXA = MOR = PYR > LEV= EPI = NOR = DA = TYR = BET = BEPH. Sample recording to each ligand is shown. Error bars represent standard error. *p<0.05; ****p<0.0001

## Supplemental Tables

**S1 Table.**
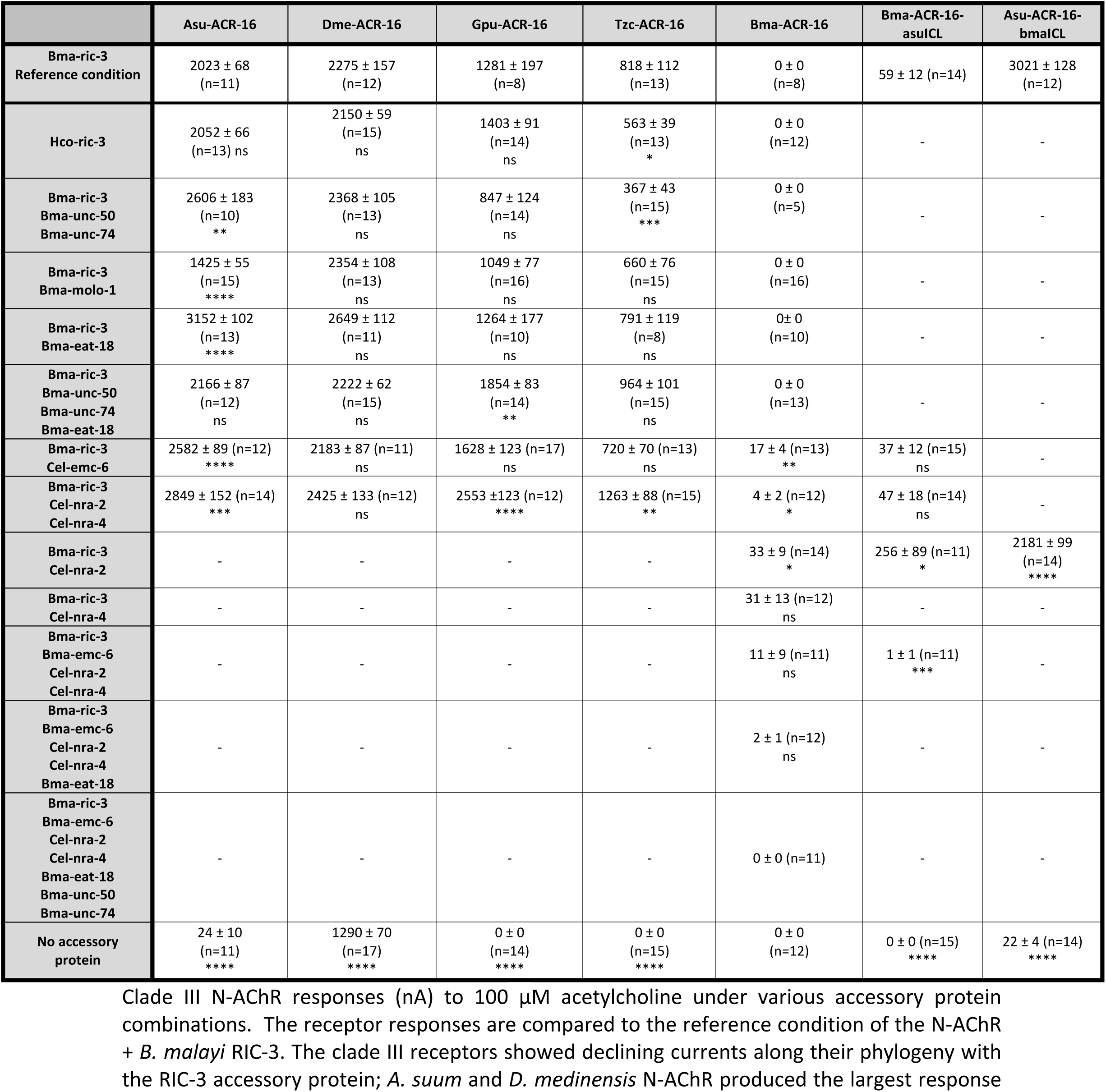
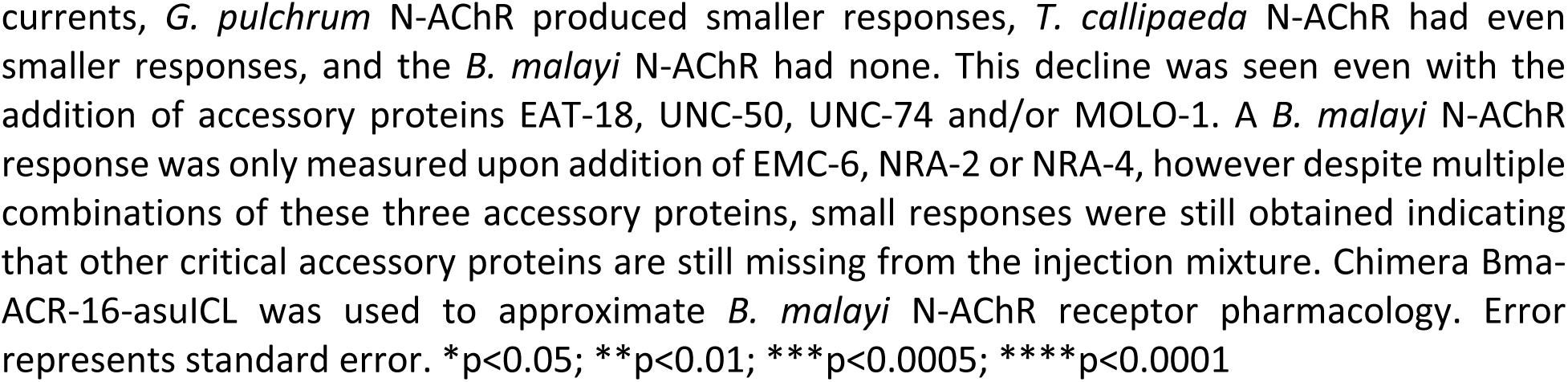
N-AChR current response with various accessory protein combinations.

**S2 Table.**
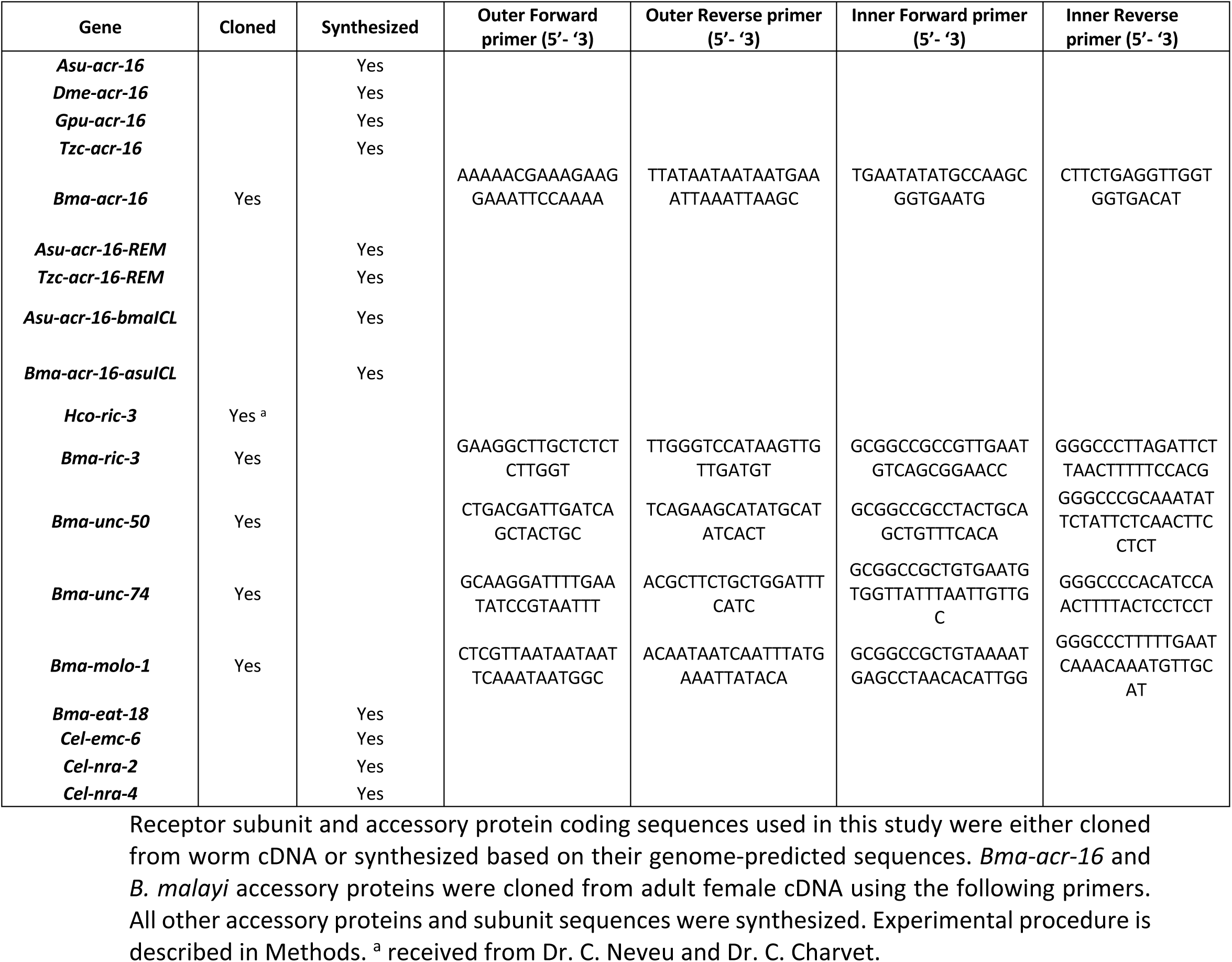
Primers used for cloning.

